# Computational modeling of an RNA-peptide world

**DOI:** 10.64898/2026.09.21.753249

**Authors:** Harutyun Sahakyan, Yuri I. Wolf, Eugene V. Koonin

**Affiliations:** Computational Biology Branch, Division of Intramural Research, National Library of Medicine, National Institutes of Health, Bethesda, MD 20894, USA

## Abstract

Emergence and evolution of functional RNA and protein structures are the central problems for understanding the origin of life. Although it is well known that catalytically active RNA elements, ribozymes, can catalyze many reactions ^1–3^, including peptide bond formation, the specifics of the transition from the hypothetical, primordial RNA world ^4–6^ to protein-based life centered at the translation system remain enigmatic ^7–9^. We developed AMES, Atomistic Molecular Evolution Simulator, and employed it to perform computer simulations of the evolution of short RNA molecules and RNA-peptide complexes. Comparison of the evolutionary trajectories and the structures of RNA molecules and RNA-peptide complexes emerging in these simulations shows that short random peptides accelerate RNA evolution, stabilize RNA folds, and boost the structural diversity of evolving RNA molecules, potentially enabling a broader range of activities. We hypothesize that the RNA world was, actually, an ‘RNA-peptide world’, in which, from the earliest stages of evolution, evolving RNA molecules interacted with short random peptides synthesized in a non-templated manner. These interactions could drive the evolution of diverse RNA structures and activities, and eventually, of the translation machinery.

---

The emergence of life remains one of the hardest problems in all of science, but parallel advances in several areas inform and constrain origin of life scenarios. Numerous studies indicate that biologically relevant molecules, such as amino acids ^10^, nucleobases and nucleotides ^11–13^ and even short peptides ^14,15^ and oligonucleotides ^16–18^ are readily produced under conditions thought to mimic those at the Hadean earth ^19,20^.

One of the cornerstones of the origin of life field is the RNA world hypothesis under which at the earliest stage of life evolution, the roles of both information carriers and catalysts were performed by RNA molecules ^4–6^. The RNA world appears to be a logical necessity because, in modern cells, nearly all the catalytic functions, in particular, genome replication, are performed by protein enzymes that are produced by the highly complex translation system which itself must have evolved in a setting where efficient replication of information carriers was already in place along with a steady supply of amino acids ^8,21^. Substantial support for the RNA world comes from the study of ribozymes, catalytically active RNAs that both occur in modern life and are designed experimentally ^1–3^. Strikingly, the key reaction of translation, the formation of the peptide bond, is catalyzed by a ribozyme which is the most highly conserved portion of the large ribosomal subunit RNA ^22,23^. A purified peptidyltransferase ribozyme as small as 64 nucleotides has been shown to catalyze non-templated peptide synthesis *in vitro* ^24,25^. Although another key step of translation, aminoacylation of tRNAs, is catalyzed by protein enzymes in all modern cells, ribozymes as tiny as pentanucleotides have been shown to catalyze aminoacylation in vitro ^26,27^. These findings strongly suggest that the translation system emerged as an RNA-based, possibly, even RNA-only molecular machine. An essential ingredient of any RNA world model is a ribozyme RNA polymerase endowed with fidelity, processivity and polymerization rate sufficient to enable efficient RNA replication and, consequently, evolution. Over the years, much effort has been dedicated to producing such an efficient ribozyme polymerases by molecular design and experimental evolution ^28,29^, and in recent breakthrough, an efficient polymerase of only 45 nucleotides (albeit joining trinucleotides rather than mononucleotides) has been obtained ^30^.

All these results are making an RNA world an increasingly realistic possibility. However, major difficulties remain. In particular, there is virtually no evidence that ribozymes can catalyze reactions of nucleotide and amino acid biosynthesis that would be required to supply the building blocks for RNA and subsequently proteins even if some amounts of these molecules can be produced abiogenically. More generally, ribozymes are inefficient catalysts compared to protein enzymes. Furthermore, there seems to be a chicken and egg paradox intrinsic to the evolution of translation: arguably, the translation machinery could not have evolved under the pressure of selection for the synthesis of proteins inasmuch as evolution has no forecast ^8^. One possible escape from this paradox is the involvement of amino acids and short peptides synthesized via non-templated processes, possibly catalyzed by ribozymes, from the earliest stages of evolution ^31,32^. Indeed, peptides can stabilize some ribozymes and act as cofactors enhancing ribozyme activity ^33,34^, and in complementary studies, short RNA molecules containing non-canonical bases have been shown to drive peptide synthesis ^32^, and amino acids have been shown to promote oligonucleotide synthesis ^35^. However, to our knowledge, interactions of RNA molecules with peptides have not been studied systematically in the context of primordial RNA evolution. If such interactions occurred in primordial systems and boosted ribozyme activities to the benefit of the respective protocells ^36^, peptidyltransferase activity of ribozymes could be selected for, potentially, kicking off the evolution of translation.

We were interested in investigating potential effects of peptides on RNA evolution through *in silico* experiments enabled by the recent advances in AI-based molecular structure prediction such as AlphaFold3 ^37,38^ and ESMfold2 ^39,40^. Previously, using this approach, we explored evolution of globular protein domains from random amino acid sequences, showing that diverse protein folds could evolve after a relatively small number of mutations, under selection for foldability and stability ^41^. In the present work, we developed AMES, Atomistic Molecular Evolution Simulator for *in silico* analysis of evolution of proteins, nucleic acids, and their complexes, providing an atom-level description of structural changes occurring during evolution. We used AMES to explore evolution of RNA folds in a simulated environment including pools of short RNA molecules and random peptides. Our simulations show that evolving RNA molecules can form complexes with peptides, resulting in acceleration of RNA evolution and diversification of the emerging structures. We surmise that evolution of RNA complexes with short peptides could jump-start the evolution of the translation system.

## Results

### Molecular evolution simulations with computed all-atom models

Deep learning-based structure prediction methods, augmented with additional techniques for accuracy evaluation, provide a good approximation of molecular interactions that define the properties of biological macromolecules which allows simulating molecular evolution from first principles, without relying on statistical information from existing sequences or other empirical data. AMES (Atomistic Molecular Evolution Simulator) combines information from all-atom structure predictions with population dynamics to imitate the evolution of proteins, RNA, and their complexes under various conditions, at the structure and the sequence levels.

AMES introduces mutations into populations of evolving sequences representing single polymer chains or complexes, such as RNA-peptide complexes in this work, and evaluates the fitness effects of these mutations to select the population members for the next generation (Figure 1A; see Methods for details). The fitness of population members is computed from their sequences, structures, and confidence metrics generated during the structure prediction (Figure 1B). Mutations include residue substitutions, indels of various lengths, duplications, permutations, and randomizations (Figure 1C). Thus, the sequence length during the simulation changes until the population stabilizes at a local minimum on the fitness landscape. New molecules can grow gradually from shorter sequences or emerge from longer ones.

**Figure 1.**
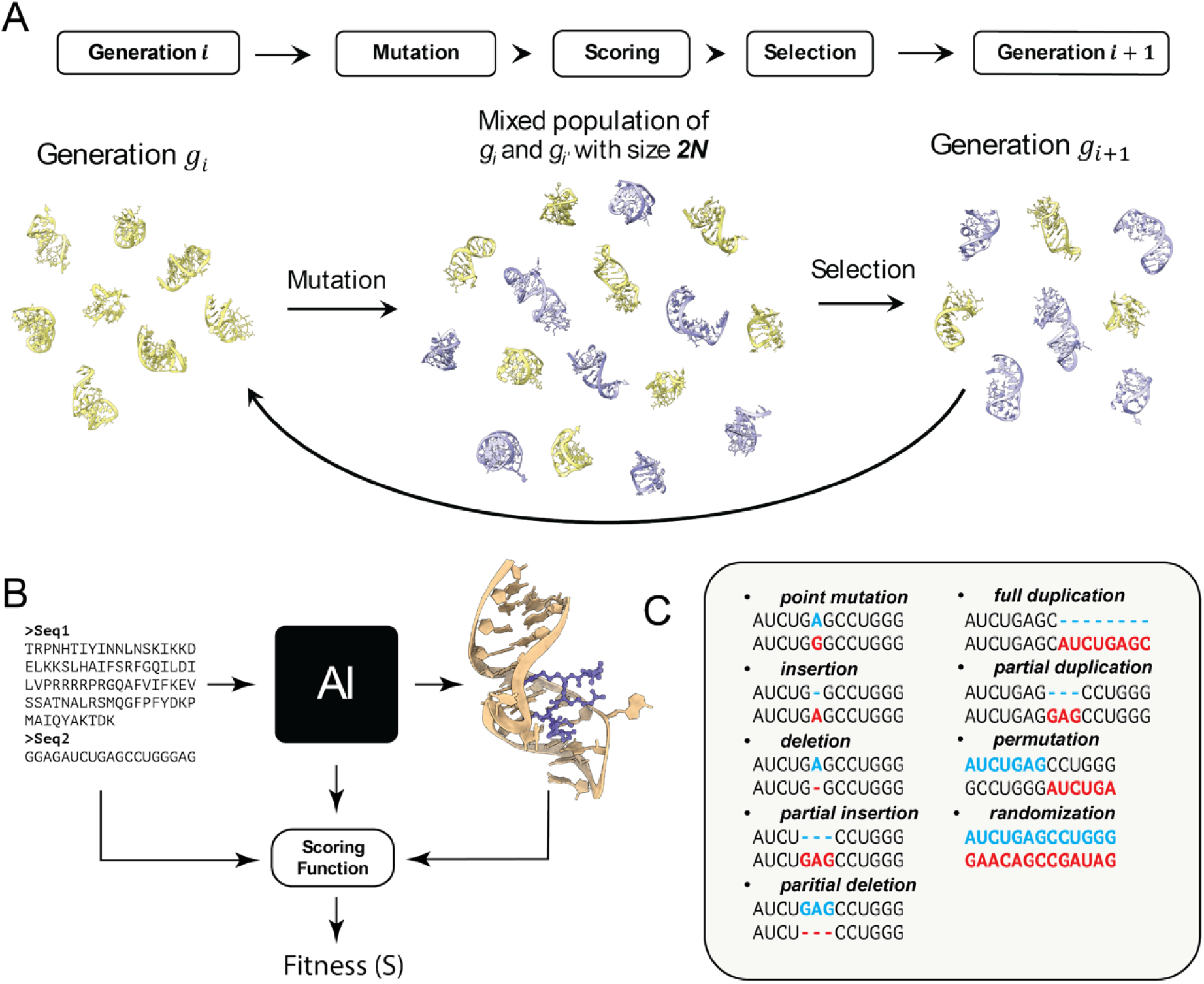
The AMES workflow. The figure shows the setup used in this study; AMES has additional options. **A)** General steps in each iteration: population of molecules mutates, creating a mix of mutated and original molecules. For all members of this mixed population, a fitness score is calculated, and selection is performed based on this fitness. This cycle repeats generation-by- generation. **B)** For calculating the fitness of a molecule or a molecular complex, first, its structure is predicted, and then the initial sequence, the predicted structure, and the confidence metrics are used to calculate the fitness score. **C)** Mutations both for amino acid and nucleotide sequences include single residue substitutions, single or multi residue indels, partial or full duplications, and circular permutations.

Given that both short peptides and oligonucleotides can be spontaneously synthesized in various conditions resembling the primordial environment, we assume that these two types of molecules would interact with each other, and these interactions could define the stability and dynamics of emerging molecular complexes. We employed AMES to investigate the emergence and evolution of small RNA folds in computer simulations of molecular evolution, where a population of RNA molecules mutated and evolved in the presence or in the absence of short, random peptides. The selective pressure and the fitness of the evolving molecules were inferred from the confidence metrics of the predicted molecular structures, namely, pTM, mean pLDDT, ipTM, and ipLDDT scores (where “i" stands for interaction; see Methods), reflecting the stability of the evolving molecules and their complexes. In these simulations, only RNA was endowed with heredity, allowing evolution via the accumulation of beneficial (along with neutral) mutations under selection pressure, whereas the peptides were randomized at each mutation step. Each simulation was initiated with a population of 100 unique, random RNA sequences, starting at 18 nucleotides (nt), with the length of evolving sequences constrained between 12 to 80 nt. In RNA-peptide evolution simulations, the RNA molecules interacted with 3-, 4- or 5-meric peptides, with both the length and the sequences of the peptides sampled randomly. For each setup, 50 independent simulations were performed, initiating the simulation with a new population of random RNA sequences each time. Trajectories from the simulations, including full lineages and the best scoring sequences from the last generations, were collected for further analysis (see Methods).

### Short random peptides change the dynamics of simulated evolution of short RNA molecules

Typical RNA folds emerging in the simulations in the absence of peptides were simple RNA hairpins, often with different types of bulges (Figure 2A, Figure S1). Although RNA hairpin is a common structural motif, searches for structural motifs in Rfam CM models and sequence searches in RNAcentral (see Methods) did not return confident matches for the *in silico* evolved RNAs. Although some similar RNA structures were detected in the PDB database in sequence- independent structural searches, these were generic, apparently non-specific similarities to hairpin structures, covering only a part of the target RNA.

**Figure 2.**
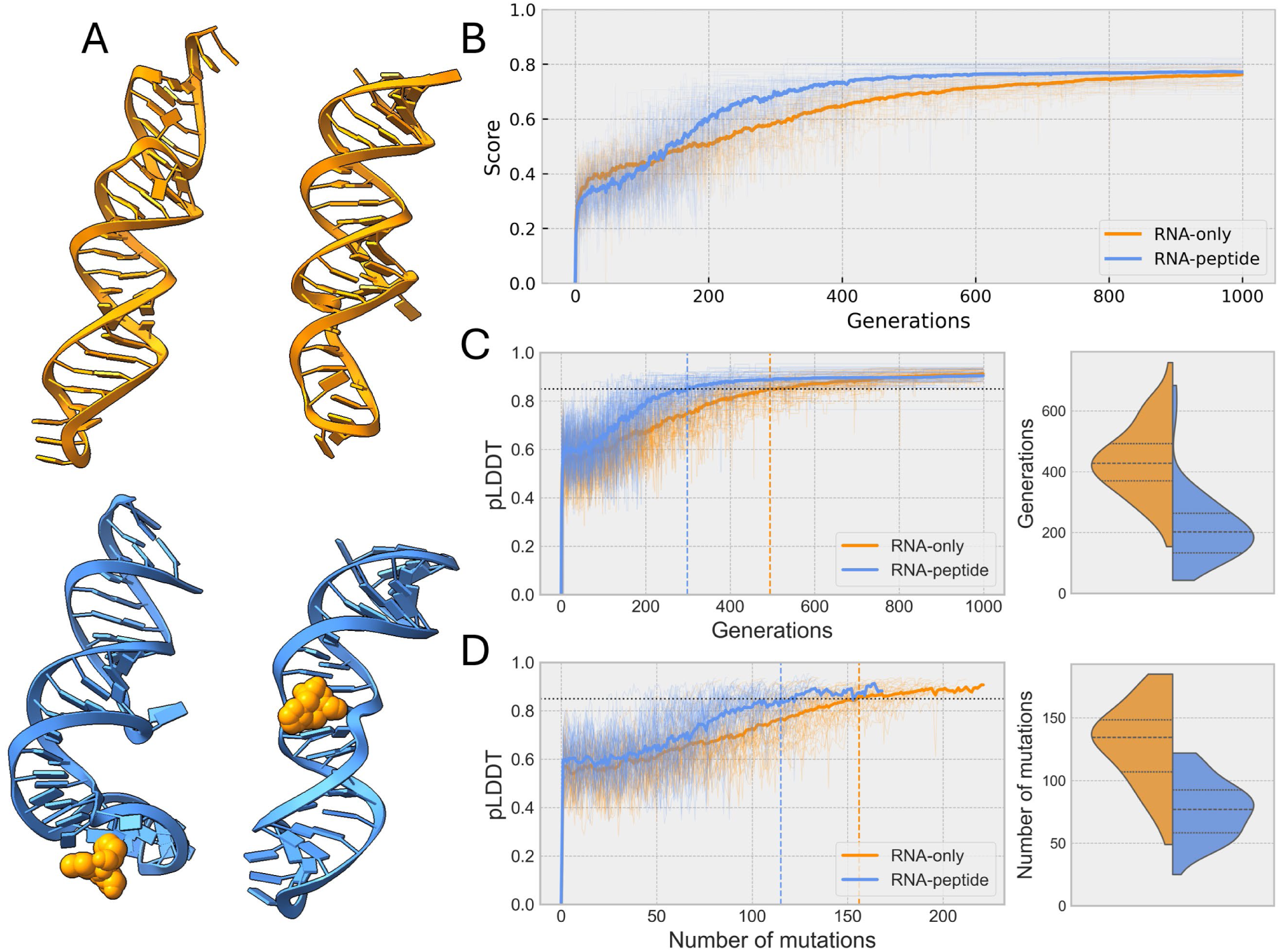
Simulation of RNA evolution in the presence and in the absence of small random peptides. **A)** Examples of RNA structures evolved in simulations with or without peptides. **B)** Mean fitness score (thick lines), and scores from individual simulations (thin, transparent lines in the background; N=50) are shown. **C)** Evolution of fold stability (pLDDT scores) in the presence and in the absence of peptides. The horizontal dashed line shows a pLDDT cutoff of 0.85. The vertical dashed lines show the number of generations when the mean pLDDT reached the cutoff. Distributions on the left show the number of generations required to reach the pLDDT score cutoff in the presence and in the absence of peptides. **D)** The number of fixed mutations required to reach the pLDDT score cut-off in the presence and in the absence of peptides, and distributions of the number of mutations required to reach the pLDDT score cutoff in the presence and in the absence of peptides. See Table 1 for more details.

In simulations with or without peptides, the fitness score increased steeply in the beginning and gradually saturated, reaching a plateau (Figure 2B). However, the shapes of the fitness curves in the absence and in the presence of peptides were markedly different. In the presence of peptides, the evolving RNA reached the fitness plateau much faster than RNA evolving in isolation; in the latter case, fitness did not completely saturate at the end of the simulation (Figure 2B). To quantify the difference in the evolution rates, we measured how many generations were required to reach a stable fold (mean pLDDT ≥ 0.85). For RNA evolving in isolation, 438 ± 120 generations (mean ± SD) were necessary to reach this threshold, whereas RNA evolving in the presence of small peptides reached it after 215 ± 116 generations (Figure 2C). Thus, interaction with peptides decreased the number of generations required to reach the cutoff, that is, in the presence of short, random peptides, the evolving RNA approached the fitness plateau about twice as fast as RNA evolving in isolation.

The number of mutations in a simulation is proportional to the number of generations, but most of these mutations are eliminated by selection, and only a small fraction of mutations that are beneficial or neutral is fixed by positive selection or drift, respectively. Therefore, we also calculated the number of fixed mutations required to reach the same threshold, which represents the length of the evolutionary lineage from the initial random sequence to the final, stable RNA fold. For RNA-only simulations, 129 ± 32 mutations were necessary to evolve a structure with pLDDT ≥ 0.85, whereas in the case of RNA evolution in the presence of random peptides, it took 77 ± 23 mutations, supporting the conclusion on the substantial acceleration of RNA structure evolution by peptides interacting with RNA (Figure 2D).

Thus, short, random peptides caused a substantial, statistically highly significant acceleration of RNA evolution *in silico* (Table 1), although the exact mechanism behind this acceleration remains unclear. Despite the similarities in the simulation protocols, including the same number of generations, population size, and selection criteria, the RNA-only simulation required many more mutations to attain a stable fold (high fitness). In this case, fitness increased smoothly while a random RNA sequence evolved into a hairpin-like fold, gradually accumulating mutations allowing the formation of Watson-Crick pairs. In contrast, evolution of RNA interacting with peptides required fixation of conformations allowing peptide binding, in addition to RNA fold stability, creating additional constraints in the search for and fixation of favorable mutations, especially, given the randomization of the peptides at each mutation step. Therefore, in the beginning of the simulations, isolated RNA gradually accumulated favorable mutations, resulting in the smooth fitness increase, whereas in RNA-peptide simulations, although in the beginning, most mutations were eliminated, after approximately 100 generations, favorable mutations started to accumulate quickly until, in most simulations, the fitness plateau was reached (Figure 2B,C).

**Table 1.**
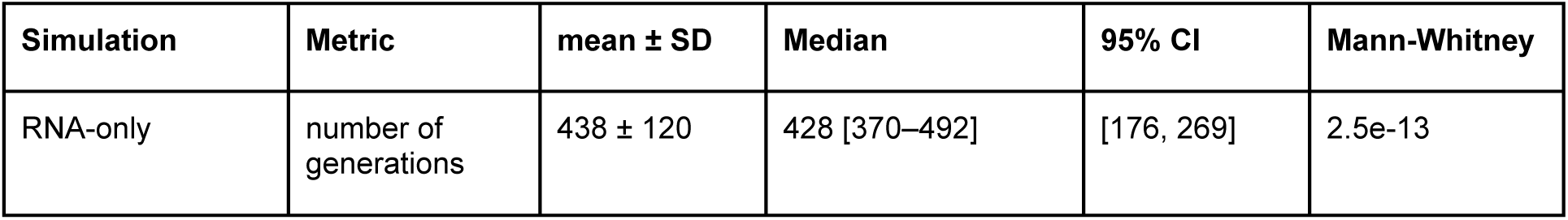
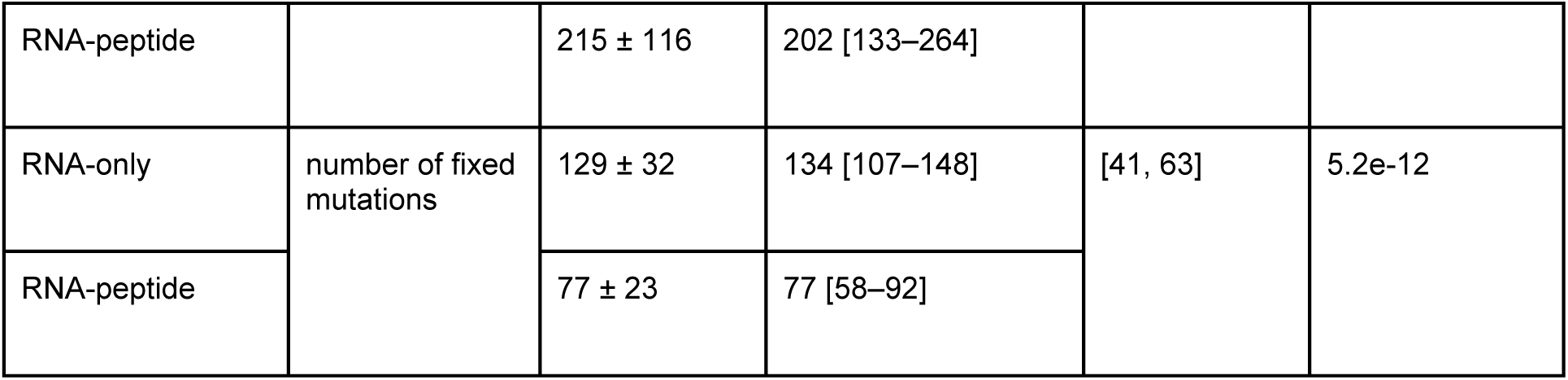
Rates of RNA evolution in the presence and in the absence of peptides.

| Simulation | Metric | mean ± SD | Median | 95% CI | Mann-Whitney |
| --- | --- | --- | --- | --- | --- |
| RNA-only | number of generations | 438 ± 120 | 428 [370–492] | [176, 269] | 2.5e-13 |
| RNA-peptide |  | 215 ± 116 | 202 [133–264] |  |  |
| RNA-only | number of fixed mutations | 129 ± 32 | 134 [107–148] | [41, 63] | 5.2e-12 |
| RNA-peptide |  | 77 ± 23 | 77 [58–92] |  |  |

The emergence of stable folds from random sequences requires the accumulation of enough stabilizing mutations to support a certain configuration of the polymer. For RNA, the simplest stable configuration is a hairpin, which we observed as the final structure in most of the simulations with RNA alone (Figure 2A, Figure 1S). When an RNA hairpin nucleates, it is unstable because only a small fraction of the bases is paired, whereas the rest of the Watson-Crick pairs are gradually formed through the accumulation of further mutations, resulting in fold stabilization. By contrast, peptides interact with unpaired nucleotides in the evolving RNA molecules, stabilizing RNA folds before the number of base-pairs becomes sufficient for stabilization of RNA alone (Figure 2A). The Root Mean Square Deviation (RMSD) of aligned RNA backbones shows that, once a stable structure emerged, it underwent only minor changes maintaining the emerged fold. The relative RMSD values of peptides calculated after aligning the RNAs showed that, after 200- 400 generations, peptides started binding the same region of the evolving RNAs (Figure 3A,B, Movie S1), despite the random generation of the peptides at each step. This observation is compatible with the emergence of a generic, non-sequence-specific peptide-binding capacity in RNAs that are exposed to peptides during their evolution.

**Figure 3.**
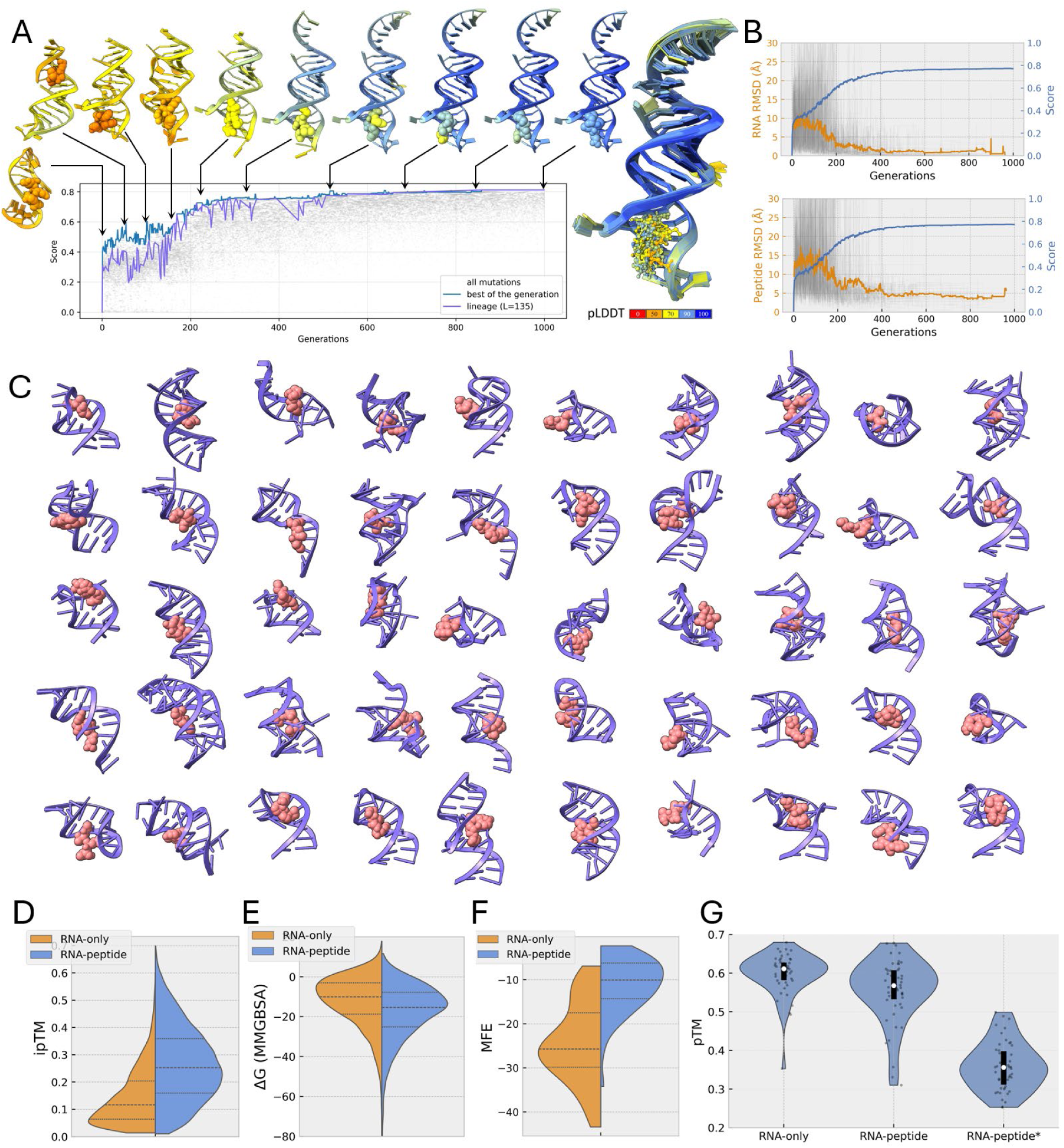
Stability of RNA folds in the presence and in the absence of peptides. **A)** An example of RNA peptide evolution dynamics. The plot shows the fitness scores for all mutations throughout the simulation (gray dots), for the mutations with the highest score in each generation (blue line) and for the lineage of fixed mutations (purple line). Structures around the plot show intermediate states with arrows pointing to the specific generation in which that intermediate state was detected. Superimposed structures starting from the 300th generation in the lineage are presented on the right. The structures are colored by the pLDDT scores. **B)** RMSD for RNA and for the peptides, measured after aligning RNA structures, with the thick line showing the mean and the thin transparent lines in the background showing individual simulations. The blue line on the second y-axis, illustrating the mean fitness score of RNA-peptide evolution is shown for comparison with RMSD. **C)** Final structures from the RNA-peptide simulations. Only RNA- peptides interaction regions are visualized **D)** ipTM score reflecting interaction of RNAs evolved in the presence and in the absence of peptides with a new set of random peptides. Split violin plot shows ipTM score distributions for each RNA set. **E)** Distribution of binding free energies calculated with MMGBSA method. **F)** Minimum Free Energy (MFE) distribution calculated from secondary structures representing the most thermodynamically stable configuration. **G)** pTM score distributions for RNA evolved in isolation, RNA-peptide complexes, and RNA from RNA- peptide complexes predicted without the peptide (denoted as RNA-peptide*, See Table 2 for statistics).

These findings suggest that bulges and non-canonical configurations in RNA structures often serve as anchors at the RNA-peptide interaction interface, as indeed observed by inspection of the binding sites where peptides interact with the RNA (Figure 3A,C). Although the peptide binding location on RNA did not show obvious, specific patterns, in most cases, the binding sites contained unpaired nucleotides (Figure 3C). Thus, RNAs evolved in the presence of peptides could be predicted to possess a greater capacity to bind random peptides than RNAs evolved in isolation. To test this prediction, we investigated the formation of complexes with random peptides for RNAs evolved in the presence and in the absence of peptides (100 new random peptides interacting with the RNAs evolved in each of the 50 simulations, that is, 5000 complexes altogether for each set of RNAs). The RNAs that evolved in the presence of peptides showed significantly higher ipTM scores reflecting the RNA-peptide complex stability than RNAs evolved in isolation (Figure 3D, Table 2, p = 7.3×10^-10^). We performed an additional calculation of RNA-peptide affinity using a physics-based approach to avoid potential bias from the AI structure prediction, which independently confirmed that RNA evolved in the presence of peptides is predisposed to random peptide binding (Figure 3E). The free energy of binding calculated using the MMGBSA method (see Methods) had a median of -11.53 kcal/mol for RNA evolved in isolation, and -16.99 kcal/mol (p = 9.1×10^-5^) for RNA evolved in the presence of peptides setup. Thus, RNAs evolved in the presence of peptides acquire a predisposition for generic, sequence-independent peptide-binding, resulting from the evolution of distinct sites in the RNA folds capable of accommodating amino acids, in contrast to hairpins in which (nearly) all bases are paired.

**Table 2.** Stability of evolved RNAs in the presence and in the absence of peptides.

| Simulation | Median MFE | Mann-Whitney | median pTM | Mann-Whitney | median pLDDT | Mann-Whitney |
| --- | --- | --- | --- | --- | --- | --- |
| RNA-only | -25.70<br>[-29.85, -17.50] | p=1.2e-10 | 0.61<br>[0.58, 0.63] | p = 0.0014<br>with RNA-peptide | 0.91<br>[0.89, 0.92] | p = 5.2e-09<br>with RNA-peptide |
| RNA-peptide* | -10.05<br>[-14.28, -6.20] |  | 0.36<br>[0.31, 0.40] | p = 7.1e-15 | 0.68<br>[0.641, 0.75] | p = 2.4e-11 |
| RNA-peptide | N/A |  | 0.57<br>[0.53, 0.61] |  | 0.85<br>[0.79, 0.88] |  |

| Simulation | Median ipTM | Mann-Whitney | MMGBSA | Mann-Whitney |
| --- | --- | --- | --- | --- |
| RNA-only | 0.13<br>[0.09, 0.2] | p=7.3e-10 | -11.53<br>[-14.24, -8.48] | p = 9.1e-05 |
| RNA-peptide | 0.27<br>[0.21, 0.3] |  | -16.99<br>[-20.05, -15.03] |  |

Together, these findings suggest that interactions with peptides allow RNA to stabilize in an alternative conformation, deviating from the energetically most favorable, near-perfect hairpin. Then, RNA evolving in the presence of peptides should be comparatively unstable in the absence of these peptides. To test this prediction, we calculated the minimum free energy (MFE) from the evolved RNA secondary structures, ignoring the peptides (Figure 3F, Table 2), which is another way to test our conjecture on the stabilizing effect of peptides, relying on a physics-based model. As predicted, RNA evolved without peptides had a substantially and significantly lower MFE with the median of -25.7 kcal/mol, indicating much higher stability, compared to RNA evolved with peptides which had a median of -10.05 kcal/mol (p = 1.2×10^-10^). A similar result was obtained by analyzing the RNA tertiary structures. We predicted and analyzed the final structures of RNA molecules evolved in the presence of peptides, with the peptides removed. Without the interacting peptides, the median pTM score dropped substantially and highly significantly, from 0.568 to 0.356, and the median pLDDT score dropped from 0.853 to 0.678 (Figure 3G, Table 2). Together, these observations clearly demonstrate the stabilizing effect of the interaction with peptides on the evolving RNA.

### Short random peptides drive diversification of emerging RNA folds

The ability of peptides to stabilize RNA folds in energetically unfavorable conformations implies that RNAs evolving in the presence of peptides should also adopt more diverse structures. To test this prediction, we performed an additional set of simulations with stricter length constraints to minimize the effect of the RNA size on the evolving structures. The RNA length was constrained to a maximum of 40 nucleotides, and separate simulations were performed for peptides of 3, 4, or 5 amino acids.

The previous observation of the faster convergence of RNA folds in the presence of peptides held in these simulations. In the presence of peptides, RNA reached the pLDDT cutoff of 0.85 much faster than RNA alone, and furthermore, this saturation was reached faster with shorter peptides (Figure S2), as could be expected given the smaller combinatorial space available for fewer amino acids. As in the previous set of simulations, in most cases, hairpin-like structures evolved (Figure 4A). RNA evolving in isolation formed stable hairpins with canonical base-pairing, with the main structural differences observed in the loop (Figure 4A). Peptides increased the diversity of the evolving RNA folds, and although most of the evolved RNAs still formed hairpins, many of these contained larger bulges distorting the hairpin fold (Figure 3C). This outcome is consistent with our previous observations that, in the presence of peptides, RNA was stabilized in a relatively high free energy conformation, instead of gradually evolving towards the most stable hairpin in which most nucleotides are base-paired. To quantify and compare the diversity of the emerging RNA folds, we clustered RNA structures evolved in the simulations using a sequence-independent structural comparison based on TM-scores. When RNA backbones were clustered with the TM-score threshold of 0.4, which is considered indicative of RNA fold similarity ^42^, the evolved isolated RNAs formed 9 clusters, and RNA evolved in the presence of peptides formed 16, 15, and 15 clusters for tri-, tetra-, and penta-peptides, respectively (Figure 4B, Table 3). The distributions of pairwise TM-scores and RMSD for RNA backbones showed the same trend, indicating a substantially greater structural diversity among the RNA folds evolving in the presence of peptides (Figure 4C, Table 3). The same diversifying effect of peptides was observed in simulations with longer RNAs (Figure S3).

**Figure 4.**
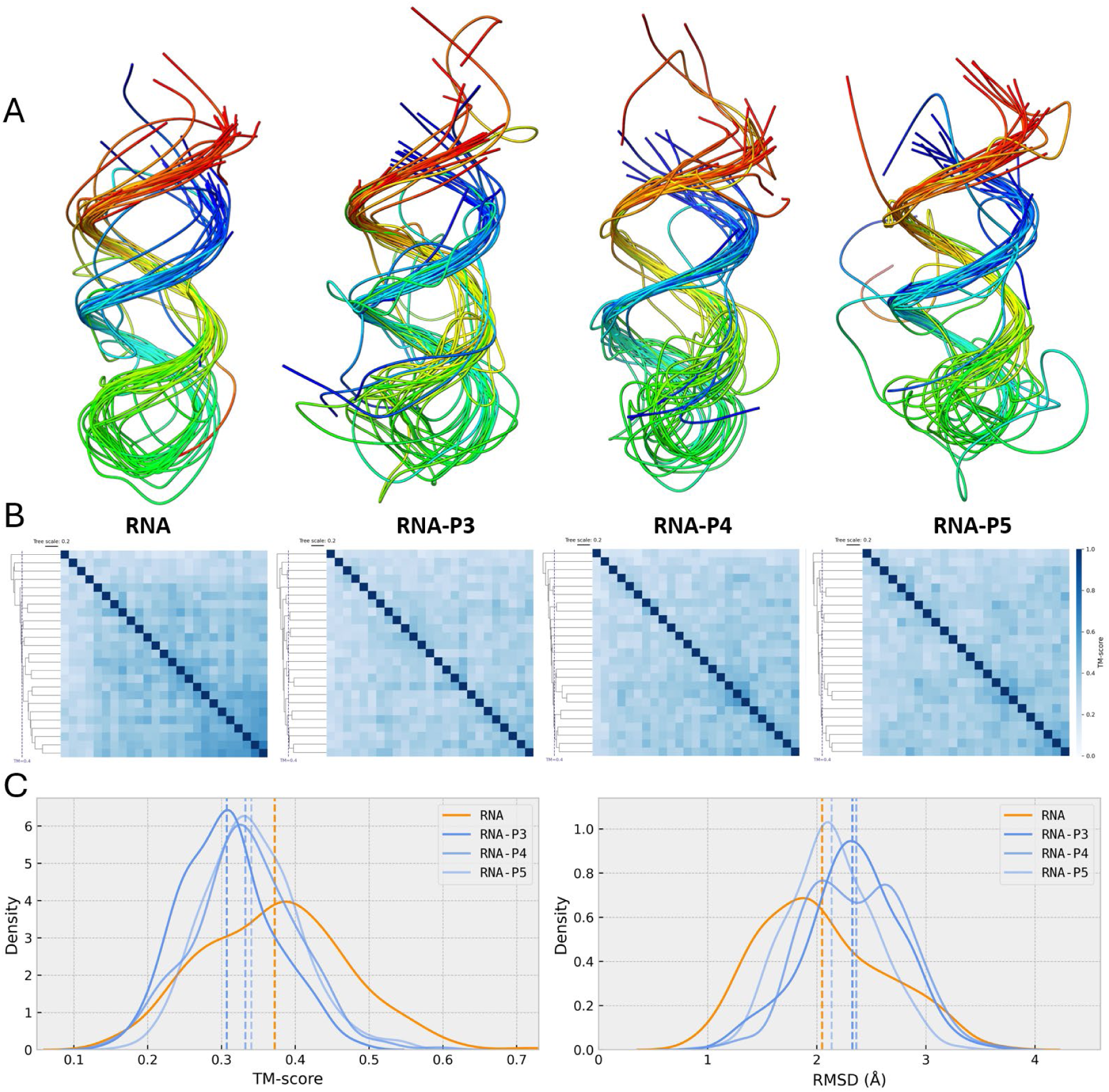
Diversity of RNA structures evolved in isolation or in the presence of peptides. **A)** Superimposed RNA backbones rainbow colored from 5’ (blue) to 3’ (red). **B)** Heatmaps showing pairwise all-vs-all comparison and UPGMA trees used for clustering. P3, P4 and P5 denote tri-, tetra- and pentapeptides, respectively. **C)** Distribution of pairwise TM-scores and RMSD with dashed lines showing medians, and labels indicating simulation setup.

**Table 3.** Clustering and pairwise comparison of RNA structures.

| simulation setup | Nr. of simulations | Nr. of clusters | Largest cluster | Mean cluster size | Nr. of singletons | Pairwise TM-score median | Pairwise RMSD median |
| --- | --- | --- | --- | --- | --- | --- | --- |
| RNA | 25 | 9 | 13 | 2.78 | 5 | 0.38<br>[0.30, 0.44] | 1.95<br>[1.60, 2.44] |
| RNA-P3 | 25 | 16 | 4 | 1.56 | 10 | 0.31<br>[0.26, 0.34] | 2.36<br>[2.08, 2.60] |
| RNA-P4 | 25 | 15 | 6 | 1.67 | 12 | 0.33<br>[0.29, 0.38] | 2.35<br>[2.02, 2.69] |
| RNA-P5 | 25 | 15 | 5 | 1.6 | 11 | 0.34<br>[0.30, 0.38] | 2.13<br>[1.88, 2.41] |

## Discussion

There is no doubt that RNA played a central role in the origin and early evolution of life. An RNA world, in which RNA molecules performed both informational and catalytic roles, appears to be an unescapable, essential early stage as suggested by the logical necessity of such a system to solve the chicken-and-egg paradox of the origin of translation and strongly supported by the presence of ribozyme relics in all modern organisms, most notably, in the peptidyltransferase center of the ribosome ^22,23^ ^24,25^. The RNA World hypothesis has been further boosted by the advances in experimental demonstration of a variety of ribozyme activities, in particular, oligonucleotide aminoacylation, ligation and polymerization ^26–30^. Nevertheless, serious challenges to the RNA world scenario remain. All the extensive study of ribozymes notwithstanding, their catalytic versatility remains limited, and in particular, there is little to no evidence of ribozyme catalysis of metabolic reactions. Furthermore, the path from the RNA world to the modern biological information system centered on the translation machinery is far from being clear. Arguably, some of these difficulties can be overcome in a RNA-peptide world where amino acids and small peptides produced by non-templated processes, possibly, aided by catalysts, including ribozymes, would be involved in the evolution of life from its earliest stages ^8,43^. This early involvement of peptides could enhance catalytic activities of ribozymes and set the stage for the origin of translation.

Here, we designed and implemented AMES, a computational engine for *in silico* exploration of macromolecular evolution and used it to investigate potential effects of peptides on the evolution of small RNAs. With the stability of RNA molecules or RNA-peptide complexes set as the target of selection, addition of short, random peptides to the evolving RNA pool dramatically changed the evolutionary trajectories. Instead of ‘boring’ hairpins that predictably evolved in simulations with RNA alone, peptides caused the emergence of far more diverse RNA structures that were comparatively unstable by themselves but were substantially stabilized by peptide-binding. Put another way, peptide-binding apparently relaxed the selection for base-pairing and diverted the evolutionary trajectory towards more complex and diverse RNA structures. RNA molecules evolved in the presence of peptides showed increased, generic, non-sequence-specific affinity for peptides. Notably, in the simulations including peptides, stable RNA-peptide complexes evolved much faster than stable RNA molecules in RNA-only simulations. These findings suggest that the earliest stage in the evolution of (pre)life was an RNA-peptide world in which peptides boosted the evolution of diverse RNA structures and hence potential activities. Furthermore, in this hypothetical RNA-peptide world, selection could favor evolution of peptidyltransferase ribozymes because protocells harboring such ribozymes would benefit from efficient production of peptides ^36^. Unlike in our previous work where about half of the in silico evolved protein folds were significantly similar to ones found in nature ^41^, here, we detected RNA structures that only generically resembled known folds, such as simple hairpins, although with very low sequence similarity. This is most likely the case because, despite the simplicity of the hairpin motifs, the space of accommodating sequences and the corresponding covariant structures is so vast that it is extremely unlikely to independently converge to a previously found solution. Nevertheless, the diverse RNA structures evolving in the presence of peptides are expected to provide fertile ground for selection of ribozymes, especially, those stabilized and/or stimulated by peptides.

The setup of our *silico* evolutionary experiments is certainly simplified and, in particular, fully relies on the assumption that efficient RNA replicases already existed in the RNA-peptide world. Provided, however, that this assumption holds, the coexistence of short RNA molecules with peptides within primordial protocells seems to be realistic, and hence so does the enhancement of RNA evolution by peptides. The complexity of the origin of life enigma is such that it cannot be solved in one clean sweep but only chipped into from different angles. In our previous work, we showed that evolution of small, globular protein domains can be relatively straightforward and fast once an efficient translation system is in place ^41^. The results of this study suggest an early RNA- peptide stage in the evolution of (pre)life and illuminate a potential early step in the evolution of the translation system.

## Methods

### Molecular evolutionary simulations

Simulations presented in this study were conducted using AMES, which was developed as a general framework for simulating molecular evolution with all-atom structural models. It allows evolutionary simulation of protein, nucleic acids, and their complexes in different conditions using sequences and corresponding structural models. Simulations are performed using population dynamics, where molecules mutate and change over time, generation by generation. A population has a fixed size *N*, where each member of the population mutates, creating an intermediate pool of size 2*N*, consisting of the generation *g* and its mutated variant *g*’ (Figure 1A). Mutations allowed in this study are illustrated in Figure 1C, although more mutation schemes are available in AMES. The mixed population is used to stochastically sample *N* members of the next generation. The probability of selecting a molecule or complex from the mixed generations *g_i_* + *g_i_*_′_ to generation *G* + 1 is calculated as

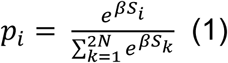

Where *S_i_* is fitness score for each molecule *i* in the population, and *β* is the inverse of the temperature controlling the selection strength.

AMES calculates fitness from different sequence and structure features combined with confidence metrics produced by a structure prediction method that serves as an oracle for predicting stability and interactions for molecules and molecular complexes (Figure 1B). In the study, *S_i_* was calculated based on confidence metrics provided by the ESMfold2 as

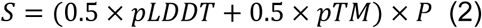

for isolated RNA molecules or as

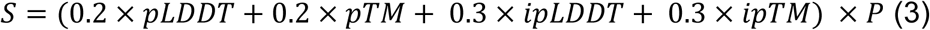

for RNA-peptide complexes. Here, *pLDDT*, *pTM* and *ipTM* are obtained directly from structure prediction and *ipLDDT* is calculated using *pLDDT* for atoms between two chains for residues located ≤ 5 Å from each other in Euclidean distances. In Eq 2 and 3, *P* is the penalty is for violating sequence minimum (*C_min_*) or maximum (*C_max_*) length constraints and for steric clashes (*SC*).

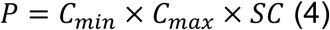

here *C_min_* and *C_max_* are

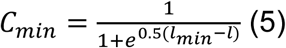

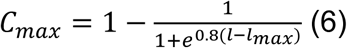

where *l* is the length of the chain and *l_min_*/*l_max_* are corresponding length constraints. Thus, length constraint scales from 0 to 1, and reduces *S* by half if the length of a constrained chain becomes *l_min_* or *l_max_*, and lower if surpasses these values. Clash score (*CS*) is calculated as

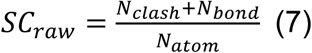

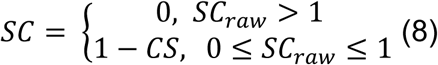

where *N_clash_* is the number of steric clashes, *N_bond_* the number of backbone covalent bond length violations, and *N_atom_* the number of atoms.

In these simulations, ESMfold2 ^40,44^ was used to predict the structures of RNAs and RNA-peptide complexes. Specifically, *esmfold2-2026-05* model with 3 diffusion samples, 3 loops, and 50 sampling steps was used. In all experiments, the simulations were run for 1000 generations, with a population size of 100, and each time, initiating the simulation with a new set of random sequences. In each generation, each RNA mutated once, and mutations included a single nucleotide substitution, a single or multiple residue indels, full or partial chain duplication, or circular permutation. The probabilities of different mutation types are provided in table S1. In simulations where peptides were present, the peptide pool was randomized with each round of RNA mutations, sampling each amino acid residue randomly with a uniform probability distribution. At the start, the RNA sequences had a length of 18, and peptides, when present, had a length of 3,4 or 5. The length of the RNA during the simulations was restricted with soft sigmoidal constraints with a minimum length of 12 and a maximum length of 40 or 80, as indicated in Results. Simulations were performed with temperature annealing by linearly increasing *β* from 1 to 10, which gradually decreased sampling temperature and increased selection strength (see Eq 1).

### RMSD calculation

RMSD of a lineage from a trajectory shows how much a structure changes during the simulation. It is calculated using USalign, after aligning each structure to its preceding structure (*i+1* to *i*) and when mutations introduce minimal changes in the structure RMSD becomes lower. The relative RMSD is calculated for the second chain (in this case for the peptide), after aligning the first chain i.e. RNA. Relative RMSD shows how much the second chain (peptide) changes it position relative to the first chain (RNA) throughout the simulation. Relative RMSD basically shows how much the peptides “jumps around” the after each mutation, which means that when relative RMSD is low, peptide binds approximately to the same site even if sequence changed completely.

### RNA secondary structure prediction

RNA secondary structures and Minimum Free Energy (MFE) were computed from RNA sequences using ViennaRNA package (*RNA.fold* with default parameters was used ^45,46^/

### Free energy of RNA-peptide binding

We used MMPBSA.py for calculating binding free energy of RNA-peptide complexes, which were initially minimized in implicit solvent with 1000 relaxation steps using openMM ^47,48^. The ff19SB and OL3 amber force fields were used for peptides and RNA respectively ^49,50^. Free energy was calculated using MMGBSA with the GBn1 model, which showed the best accuracy for protein-RNA complexes ^51^.

### Clustering of RNA structures

All-vs-all comparison was performed with avaclust (commit 07b1c82), using US-align ^42^ for aligning RNA structures and clustering the UPGMA tree with a 0.4 TM-score cutoff (*avaclust -i pdb/ -o clust/ --cutoff 0.4 --chain A*).

### Statistics

For comparing samples and calculating significance, Mann-Whitney U test implemented in SciPy was used (*scipy.stats.mannwhitneyu*). To calculate significance of ipTM differences illustrated on Figure 2D, we averaged ipTM score for each RNA interacting with 100 random peptides by taking the mean ipTM from 100 different peptides interacting with the same RNA.

### Visualization

ChimeraX ^52^ and AMESViewer, which is part of the AMES package, were used for visualizing structures and evolutionary trajectories.

### Sequence and structure search

RNA sequences were searched against Rfam ^53^ using cmscan ^54^ (sequence vs CM models), sequences in RNAcentral ^55^ were searched using nhmmer ^56^. RNA structures in PDB ^57^ containing less than 3000 residues, were searched with US-align.

## Data and code availability

AMES is available at https://github.com/sahakyanhk/ames, AMESViewer at https://github.com/sahakyanhk/amesviewer, and avaclust at https://github.com/sahakyanhk/avaclust. The results of evolutionary simulations, extracted lineages, and analyses, including clustering, structure, and sequence searches, are available at https://zenodo.org/records/22859083

## Author contributions

H.S. and E.V.K. incepted the project; H.S. performed research; H.S., Y.I.W. and E.V.K. analyzed the data; H.S. and E.V.K. wrote the manuscript that was edited and approved by all authors.

## Acknowledgements

E.V.K. is grateful to Puri Lopez-Garcia for stimulating discussions. H.S., Y.I.W., and E.V.K. are supported by the Intramural Research Program of the National Institutes of Health (NIH). This work utilized the computational resources of the NIH high-performance computing (HPC) Biowulf cluster (https://hpc.nih.gov). The contributions of the NIH author(s) are considered Works of the United States Government. The findings and conclusions presented in this paper are those of the author(s) and do not necessarily reflect the views of the NIH or the U.S. Department of Health and Human Services.

