## Supplemental figures and tables for "Computational modeling of an RNA-peptide world"

### Supplementary materials

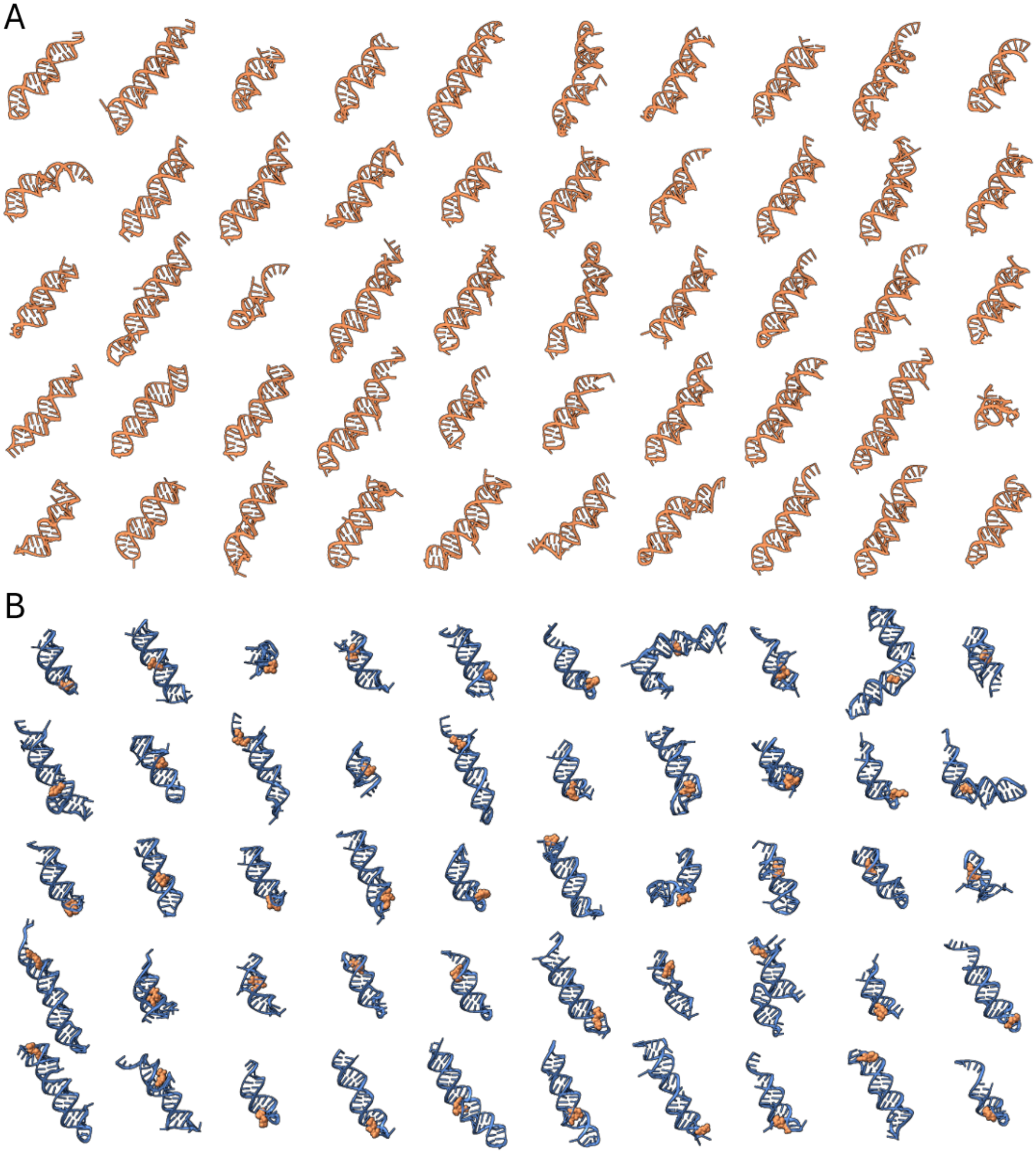


**Figure S1.** RNA structures evolved in simulations. **A)** RNA-only, **B)** RNA-peptide.

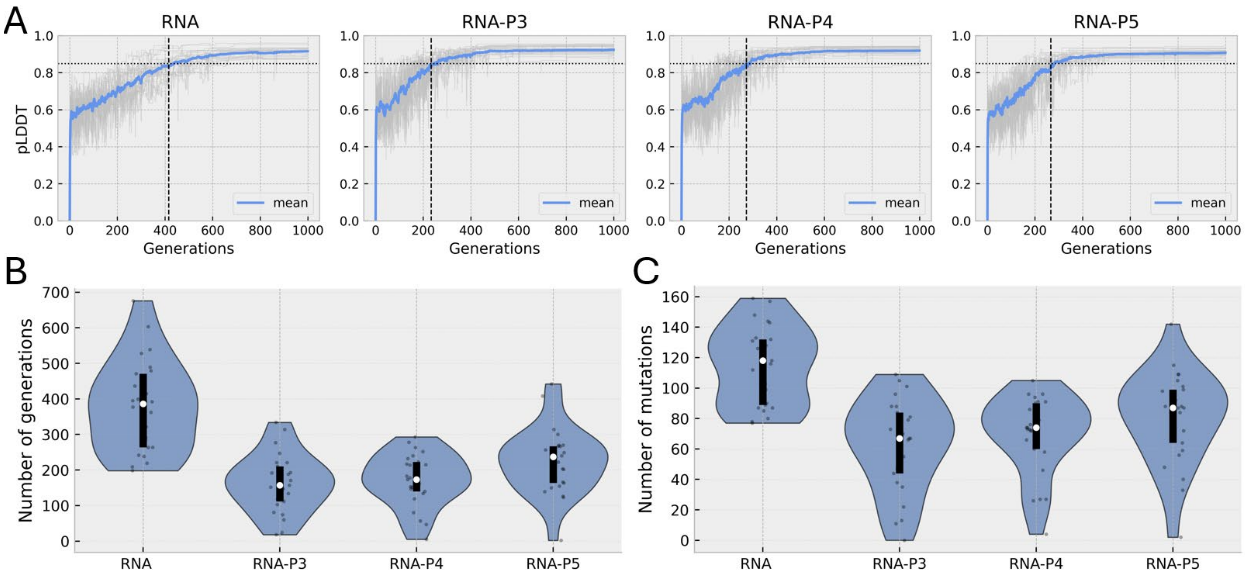


**Figure S2.** Mean pLDDT of RNA evolving in isolation and peptides. **A)** Mean pLDDT averaged across simulations and individual simulations is shown as gray lines in the background. The horizontal dashed line shows a pLDDT cutoff of 0.85. The vertical dashed lines show when the mean pLDDT reached the cutoff. **B)** Distributions showing the number of generations necessary for the individual simulations to reach the cutoff. **C)** The same for the number of mutations. N=25.

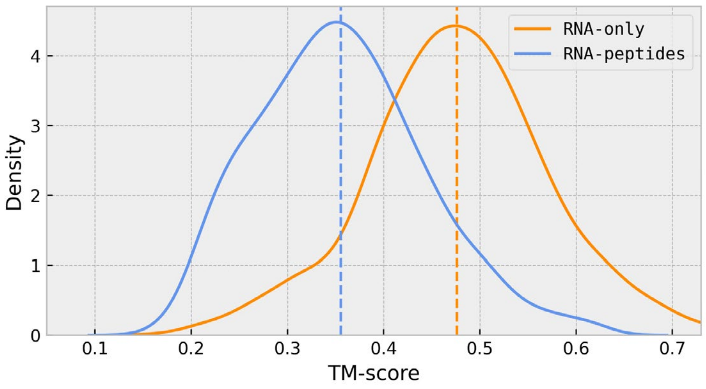


**Figure S3.** Pairwise TM-score distribution for RNA-only and RNA-peptide structures illustrated in Figures 1A and S1.

**Table S1. Frequencies of different mutation types**.

| Mutation types | Probability |
|---|---|
| Adenine | 0.127 |
| Uracil | 0.127 |
| Guanine | 0.127 |
| Cytosine | 0.127 |
| Single residue insertion | 0.127 |
| Single residue deletion | 0.127 |
| Partial insertion | 0.051 |
| Partial deletion | 0.115 |
| Circular permutation | 0.013 |
| Partial duplication | 0.051 |
| Full duplication | 0.006 |
| Randomization | 1* |

*applied to peptides independently

**Movie S1.** Example of RNA-peptide evolution. RNA with a maximum length constraint of 80 nt and peptides with 3, 4, or 5 amino acids sampled randomly (RNA-peptide, run049).
